# Vibrational optoacoustic detection of lipid-membrane dynamics enables label-free imaging of cell membrane potential

**DOI:** 10.64898/2026.06.18.733116

**Authors:** Francesca Gasparin, Jiawang Qiu, Ádám Apró, Ingo Burtscher, Heiko Lickert, Vasilis Ntziachristos, Miguel A. Pleitez

## Abstract

Current technologies to monitor membrane potential are either highly invasive and perturb the integrity of the membrane, use labels that compromise biological behavior, or are limited in sensitivity and do not enable simultaneous monitoring of multiple cells and cell populations. Here, we present Mid-IR Assessment of Conformation in Lipids by Ensemble Sensing (MIRACLES) that, by detection of molecular vibration of lipid acyl chains under cell-membrane’s electric field dynamics, achieves highly sensitive label-free imaging of membrane potential dynamics in living cells. MIRACLES leverages lipid conformational changes within the plasma membrane as intrinsic indicator for cell membrane depolarization and hyperpolarization. As proof-of-concept, we apply MIRACLES to monitor membrane depolarization during glucose stimulated insulin secretion in β-cells at single-cell level and achieve assessment of β-cell functionality in real time. These results highlight the potential of mid-IR optoacoustic as a powerful tool for indirect, label-free potential assessment of cellular metabolic activities.

## Introduction

Cell membrane potential (MP) play a key role in essential physiological functions such as: cell-cell communication in neurons, contraction and relaxation of muscle cells, release of insulin in pancreatic beta (β)-cells,^1^ etc. For instance, insulin secretion is tightly coupled to precise oscillations in membrane excitability, which are considered an indirect parameter to assess diabetes pathophysiology and β-cells functional resilience.^2–5^ Indeed, detection of MP is often used as an indirect hallmark to track the efficiency of cellular activities, providing electrophysiological insights that are inaccessible from downstream readouts alone. However, despite the importance of MP, current methods present significant challenges for monitoring membrane voltage in multiple cells. For instance, patch-clamp electrophysiology, the most common technique to measure MP, offers high sensitivity and temporal resolution, but at the expenses of disrupting integrity of cell membrane and allowing only monitoring of one cell at a time.^6^ Microelectrode arrays (MEA) allow for the monitoring of multiple cells at a time,^7^ however their implantation is not biocompatible and can cause cellular loss.^8^

Alternatively, imaging modalities such as fluorescence microscopy using voltage-sensitive dyes (VSDs) and genetically encoded voltage indicators also allow imaging MP changes in multiple cells simultaneously. Nevertheless, when applying extrinsic fluorescence indicators, the readouts are affected by artifacts arising from photo-bleaching, membrane perturbation, or poor biodistribution of the labels.^9,10^ Although fluorescence lifetime imaging (FLIM) alleviates the limitations connected to photo-bleaching and has shown to be able to image MP in thousands of cells with an accuracy superior to patch-clamp by 10-23 mV, and an accuracy eight times better than conventional fluorescence intensity microscopy, FLIM measurements can be affected by external factors such as temperature or ionic strength.^11^ Photoacoustic imaging of membrane potential using the voltage-sensitive probe dipicrylamine (DPA) reported fractional signal increases of >40% at ΔV = 100 mV and demonstrated imaging beyond the optical diffusion limit in mouse brain; however, it requires the loading of exogenous voltage probes, baring the same limitations of VSDs.^12^ As an alternative to patch-clamp or exogenous labels, full-field interferometric imaging showed label-free, non-invasive detection of action potentials in human embryonic kidney (HEK) cells by recording their deformation down to approximately 3 nm. Despite its label-free/non-destructive advantage, however, full-field interferometric imaging requires long averaging times and consequently reduces imaging speed which precludes the detection of fast MP dynamics.^13^ Additionally, second harmonic generation (SHG) in combination with wide-field illumination has been able to image the MP of mammalian neurons label-free and with a millisecond time scale. Nevertheless, the signal obtained by the alignment of water with lipids is weak (SNR ∼4 at 100 ms acquisition rate) and is influenced by signal contributions from the microtubule in the cytoskeleton and membrane lipid asymmetry, which are unrelated to MP and need to be compensated.^14^

Recently, within the vibrational microscopy techniques, hyperspectral stimulated Raman scattering (hSRS) combined with voltage-clamp has allowed detection of the MP of neurons by monitoring the CH_3_ vibrations of proteins at 2930 cm^-1^ (Fermi resonance peak). Single-mammalian neurons were monitored at high-speed and with subcellular resolution, inducing large MP changes (∼100 mV) with patch-clamp devices, thus making uncertain whether hSRS can monitor physiological alterations to MP independently.^15^ Conclusively, to date, no method has been proven to effectively detect MP in multiple cells in a non-destructive and label-free manner, preserving cell viability for longitudinal monitoring and downstream functional assays. We hypothesized that the high chemical specificity and sensitivity of live-cell mid-infrared (mid-IR) optoacoustic microscopy^16^ to detect molecular conformation^17^ and lipid content^18^ together with the action of transmembrane voltage on lipid membrane dynamics^19^ can enable label-free, non-destructive monitoring of MP in multiple cells.

To demonstrate this hypothesis, we developed Mid-IR Assessment of Conformation in Lipids by Ensemble Sensing (MIRACLES), a vibrational optoacoustic method for detecting lipid acyl chain dynamics under the action of cell’s transmembrane electric field. MIRACLES’ working principle was first demonstrated in controlled depolarization and hyperpolarization experiments on human embryonic kidney cells (HEK293) and subsequently applied to monitor MP dynamics in rat and mouse insulinoma β-cells (INS1 and MIN6, respectively) during glucose-stimulated insulin secretion (GSIS) experiments. MIRACLES sensed changes in lipid dynamic of β-cells upon uptake of glucose, revealing a progressive increase in vibrational optoacoustic signal that correlates with the depolarization associated with insulin release across the stages of GSIS. Beyond functional assessment of β-cells, MIRACLES is proposed here as an alternative to fluorescent or electrophysiological approaches of membrane potential detection that avoids phototoxicity and replace invasive cellular disruption. We anticipate that MIRACLES will enable label-free imaging of MP across a wide range of biological contexts, for instance in neuron action potential, muscle contraction and cardiac myocyte dynamics, without perturbing biological function. By providing access to intrinsic functional electrophysiological information, not accessible when probing only downstream events, MIRACLES opens a new avenue for interrogating cellular activity.

## Results

### Mechanism for endogenous contrast of membrane potential

Despite its fundamental role in many physiological processes, the impact of MP perturbations on the plasma membrane structure and dynamics remains poorly understood. Therefore, we first investigated the basic mechanism for intrinsic MP contrast generation resulting from changes in the plasma membrane voltage. **Fig. 1a** illustrates the two potential molecular mechanisms underlying MP according to the literature: (i) orientational, which causes a conformational transition of the phospholipid chain,^20–22^ or (ii) electronic, which induces a vibrational Stark effect (VSE).^19^ Conformational transitions are followed by a reorientation of the phospholipids that increases the entropy of the structural organization of the membrane, resulting in changes of the orientation angles (θ) between C-H bonds of the acyl chain and the θ of the P-N vector of the lipid head group. As a consequence kinks of the double bonds between carbon 9 and 10 in the acyl chain have been observed.^20–22^ Changes in the membrane conformation generated by variation of the potential cause an increase in the absorption coefficient of the CH_2_ phospholipid signal at 2852 cm^-1^ (ΔI).^23^ Differently, the VSE theory implies a change in the electric dipole of the phospholipid molecules. When an electric field is applied, the phospholipids align their dipole with the electric field shifting their vibrational transition energy levels from ground to excited state. This shift between the vibrational states generates a vibrational frequency shift (Δλ) of the lipid absorption band towards higher energy and lower wavenumbers (red frequency).^19,22^

**Figure 1.**
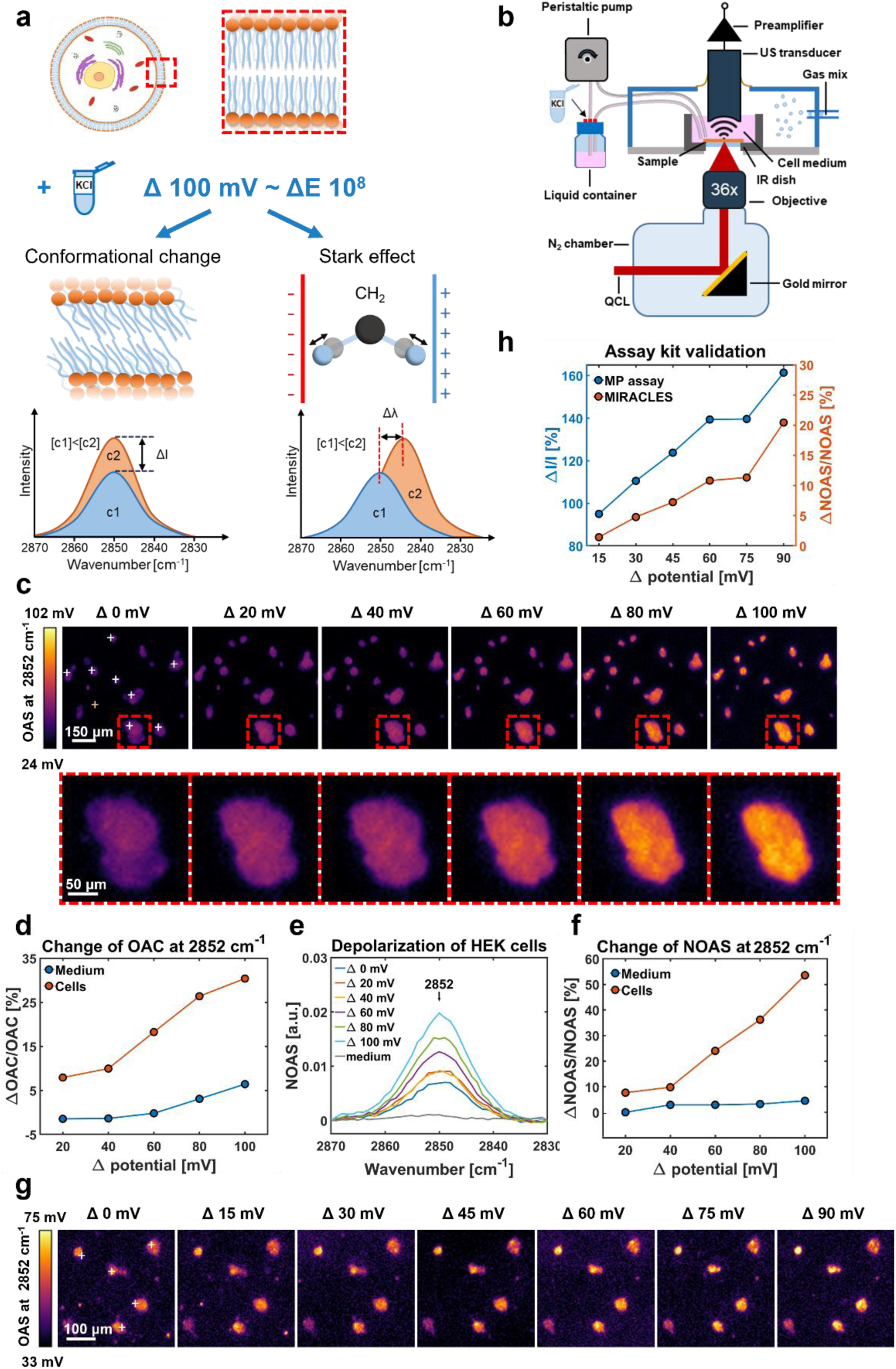
Detection of membrane potential (MP) changes in human embryonic kidney (HEK) cells using mid-infrared assessment of conformation in lipids by ensemble sensing (MIRACLES). **a)** Schematic illustration of the induction of MP changes by application of an electric field. The alteration of MP can cause either a conformational change (left) in the angle between the phospholipid head group and the acyl chain, resulting in an increase in the optoacoustic signal (OAS), or the Stark effect (right), resulting in a shift in the vibrational frequency of the OAS. **b)** Schematic diagram of MIRACLES with an integrated pump system for precise liquid flow control. A tunable quantum cascade laser (QCL) provides the excitation for the optoacoustic (OA) imaging of the cells. To generate MP changes, cells are feed with cell medium containing different concentrations of potassium chloride (KCl) using the high precision peristaltic pump. **c)** Top row: micrographs series of HEK cells acquired at 2852 cm^-1^ during the increase of MP. Bottom row: zoomed in micrographs of a single cluster of cells. **d)** Line plots showing the relative change of the optoacoustic contrast (ΔOAC/OAC) obtained for HEK cells and cell medium in (c). **e)** Normalized mean spectra of HEK cells in (c) (white crosses) and medium spectrum (yellow cross) measured at different Δ potential values. **f)** Line plots showing the relative change of the normalized optoacoustic signal (ΔNOAS/NOAS) at 2852 cm^-1^ for HEK cells and cell medium in (d). **g)** Series of micrographs of HEK cells acquired at 2852 cm^-1^ while increasing the Δ potential values. **h)** Line plots comparing the fluorescence intensity signals (ΔI/I) and the relative change in NOAS (%ΔNOAS/NOAS) at 2852 cm^-1^ when measuring MP changes in HEK cells using a microplate reader and MIRACLES, respectively.

To induce controlled and precise changes in MP in living cells, we equipped MIRACLES with a peristaltic pump modulating the concentration of KCl in the cell medium, see **Fig. 1b**. Variations of KCl concentration in the cell medium affect the resting MP (the difference of electrical potential across the membrane when cells are not transmitting signals). An increase in the extracellular concentration of K^+^ ions ([K^+^]) drives the influx of K^+^ ions into the cells to balance the concentration gradient, shifting the resting MP to more positive values and thereby causing depolarization. Conversely, a decrease in the extracellular [K^+^] causes K^+^ ions to diffuse outward, making the resting MP more negative and leading to hyperpolarization of the membrane. **Fig. 1c** shows micrographs of HEK cells acquired at 2852 cm^-1^, absorption band attributed to the symmetric stretching of the CH_2_ bond in the phospholipid acyl chain. HEK cells were imaged at different depolarization states with a variation (ΔV) in MP from ΔV=0 mV to ΔV=100 mV, induced by a stepwise increase of KCl in the medium. To gain a better understanding of how the optoacoustic contrast (OAC) is affected by the depolarization process, we calculated the relative change in OAC (% ΔOAC/OAC) in cells and medium (see **Methods**). As illustrated in **Fig. 1d**, there was an increase in OAC of more than 20% in the cells (red line), while the change in the cell medium was below 5% (blue line). The effect of depolarization on the optoacoustic signal (OAS) of the phospholipids in the membrane was also monitored by acquiring spectra within the C-H stretching region (2930 - 2770 cm^-1^). **Fig. 1e** shows the mean normalized spectra obtained from 10 clusters of HEK cells highlighted by white crosses in **Fig. 1c**, (raw spectra of cells and medium are reported in **Supplementary Fig. 1a, b**). Depolarization of the cell membrane was detected as an increase in the normalized HEK cell optoacoustic signal (NOAS) for the 2852 cm^-1^ absorption band. The relative change of HEK cells at 2852 cm^-1^ increased by 50% (red line, in **Fig. 1f**), as calculated by % ΔNOAS/NOAS (see **Methods** for details). In contrast, the relative change of the medium did show only negligible variations (blue line in **Fig. 1f**). Furthermore, we calculated the change of the area under the curve (AUC) of the mean normalized cells spectra at each voltage (see **Supplementary Fig. 1c**). The AUC increased as ΔV increased, reaching a value three times higher at ΔV=100 mV than at the resting potential (ΔV=0 mV). Contrary, the AUC of the mean normalized spectrum of the medium remained stable at different ΔV.

In addition, we induced hyperpolarization from ΔV=100 mV to ΔV=0 mV stimulating HEK cells by decreasing stepwise the concentration of KCl in the cell medium. The mean normalized spectra in **Supplementary Fig. 2a** showed a decrease of the NOAS at 2852 cm^-1^ of 100% in HEK cells (red line in **Supplementary Fig. 2b**), while the relative change in the medium was negligible (blue line in **Supplementary Fig. 2b**). The decrease in NOAS during the hyperpolarization process was also confirmed by a fourfold reduction in the AUC value (**Supplementary Fig. 2c**). The ability to detect changes in MP was examined further by observing the OAC of HEK cells extracted from micrographs obtained at 2852 cm^-1^ during hyperpolarization followed by depolarization. In **Supplementary Fig. 2d**, we first observed a 10% decrease in OA contrast when hyperpolarizing HEK cells from ΔV=60 mV to 0 mV. We then detected an 18% increase in OAC (calculated as %ΔOAC/OAC) when depolarizing the same HEK cells from ΔV=0 mV to 60 mV.

According to the analysis of the spectra and micrographs of HEK cells acquired by MIRACLES, changes in MP generated an increase or decrease in the OA signal at 2852 cm^-1^, rather than a shift in the spectral band. Thus, our observations indicate that the basic mechanism of intrinsic MP contrast in MIRACLES comes from the electric field inducing a conformational change in the phospholipid orientation.

Furthermore, to confirm that OAS/OAC changes detected by MIRACLES at 2852 cm^-1^ are caused by changes in MP, we validated our studies using an MP assay kit. HEK cells from the same tissue culture were split into two separate samples before stimulation with KCl: one dish was measured by MIRACLES (see micrographs in **Fig. 1g**), while the 96 well plate by a multi-well spectrometer. HEK cells were pretreated with an MP dye (see **Methods** for details and **Supplementary Fig. 3**), and the spectrometer measured the red fluorescence emitted by the MP dye in response to the change in MP induced by the stimulation with KCl. **Fig. 1h** shows the correlation between the emitted fluorescence signal of the MP dye (calculated as % of ΔI/I), and the OAS detected by MIRACLES (calculated as % ΔNOAS/NOAS). The similarity between the plot of the fluorescence intensity signal and the NOAS confirmed that MIRACLES is a suitable method to detect changes in MP.

Beyond unidirectional depolarization and hyperpolarization experiments using intrinsic membrane contrast at 2852 cm^-1^, a to demonstrate reproducibility, we further investigated MIRACLES capability to detect intermittent changes in MP. To this purpose, in a first experiment, MP of HEK cells was depolarized and hyperpolarized intermittently between ΔV=0 mV and 80 mV every five minutes for four times (see **Methods** for details). In **Supplementary Fig. 4a, b**, the micrographs obtained at 2852 cm^-1^ show that relative change in OAC of the cells was oscillating from low to high values (between ca. 0% and 10% increase) following the modulation in MP, while the relative change of the medium was lower than 4% (see **Supplementary Fig. 4b**). In a second experiment, serving as control, we proved MIRACLES sensitivity in detecting MP in another non-excitable cell line. The MP of HeLa cells was modulated with a stepwise elevated concentration of KCl from ΔV=0 mV to 100 mV in the cell medium, alternated by a medium with null concentration (see **Supplementary Fig. 4c**). **Supplementary Fig. 4d** shows the relative change of the OAC obtained from four cells in **Supplementary Fig. 4c** whose values were oscillating from low to high every time the medium was exchanged from null to high concentrations of KCl. Also in this case, the relative change of the OAC in the medium was negligible during the process (see blue line in **Supplementary Fig. 4d**).

### Detection of Glucose Stimulated Insulin Secretion (GSIS)

Having demonstrated the ability to detect MP changes, we next applied MIRACLES to monitor depolarization of INS1 and MIN6 during the GSIS process.^24^ **Fig. 2a** illustrates the mechanistic pathway governing insulin secretion. Rising extracellular glucose levels promote intracellular uptake, leading to increased adenosine triphosphate (ATP) production and subsequent closure of potassium (K^+^) channels. This closure induces membrane depolarization, which in turn opens calcium (Ca^2+^) channels. Ultimately the resulting influx of Ca^2+^ ions triggers insulin release. Here, we hypothesized that we can indirectly observe the release of insulin by detecting the depolarization of MP caused by the closing of K^+^ channels. The sequence of electrical and biochemical events following GSIS process makes β-cells an ideal system to demonstrate whether MIRACLES can resolve MP-linked lipid remodeling under physiologically relevant conditions.

**Figure 2.**
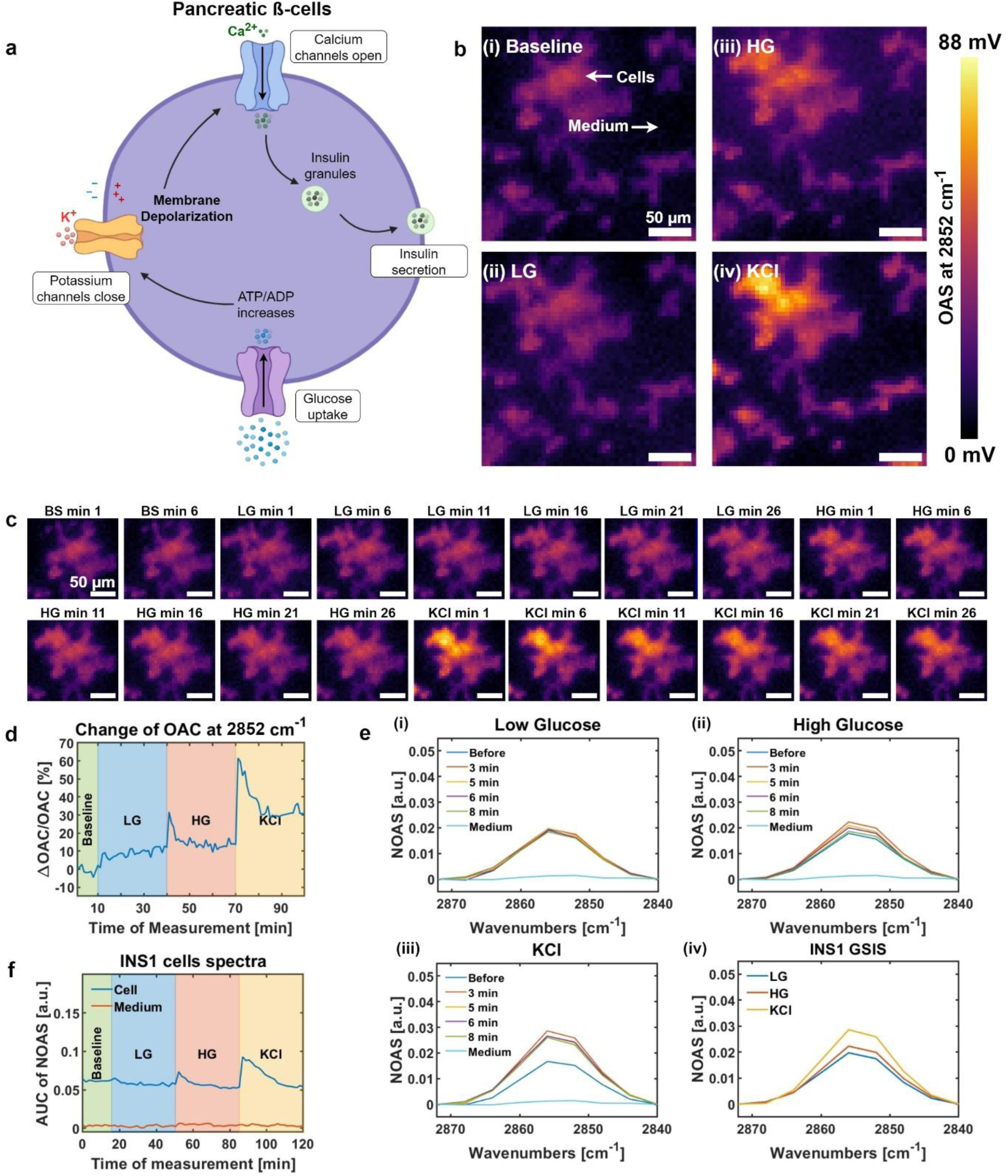
Monitoring glucose stimulated insulin secretion (GSIS) using mid-infrared assessment of conformation in lipids by ensemble sensing (MIRACLES). **a)** Diagram of the GSIS process. Glucose enters the cell causing the adenosine triphosphate (ATP) / adenosine diphosphate (ADP) levels to rise. Consequently, ATP sensitive potassium channels close causing membrane depolarization and inducing secretion of insulin. **b)** Representative micrographs of rat insulinoma (INS1) cells acquired at 2852 cm^-1^ while changing glucose and potassium chloride (KCl) concentrations in the cell medium according to the GSIS process. **c)** Series of zoomed-in micrographs of INS1 clusters showing the change in optoacoustic signal (OAS) acquired during the GSIS process. **d)** Line plot showing the relative change of the optoacoustic contrast (ΔOAC/OAC) of cells and cell medium calculated from the micrographs in **b)**. **e)** Normalized optoacoustic spectra (NOAS) acquired from single points in INS1 cell and cell medium during the different phases of the GSIS process: (i) low glucose (LG), (ii) high glucose (HG), and (iii) KCl concentrations, (iv) NOAS spectra summarizing the different phases of the GSIS process. **f)** Line plot of the area under the curve (AUC) of the spectra in (e). The increase of the AUC value after the HG and KCl phases suggests membrane depolarization. N=2.

In a first experiment, INS1 cells were subjected to sequential changes in culture medium (see **Methods** for details): (i) baseline (BS), (ii) low glucose (LG), (iii) high glucose (HG), and (iv) high KCl (KCl).^25^ To monitor changes in OAC caused by membrane depolarization, micrographs of INS1 clusters were continuously acquired at 2852 cm^-1^ over 100 minutes (10 minutes for BS and 30 minutes for each condition), at a frame rate of one image per minute. **Fig. 2b** shows representative micrographs of the different phases of the GSIS process. To better visualize the changes in the OAC as the cell medium content was exchanged, **Fig. 2c** reports a series of zoomed-in micrographs taken at 5-minute intervals. From the images in **Fig. 2b, c**, an increase of the OAC of the cells was observed when we increased the concentrations of glucose or KCl in the medium. The increase of OAC over time is depicted by the line plot in **Fig. 2d**, which reports the relative change in OAC (% ΔOAC/OAC) of cells and cell medium. In detail, the line plot highlights voltage peaks upon addition of HG and KCl to the medium (see **Fig. 2d**), indicating a rapid increase in MP consistent with the membrane depolarization following glucose and KCl stimulation.

Furthermore, depolarization of INS1 cells was confirmed by acquisition of OA spectra during the four different phases of the GSIS process. The OA spectra were acquired in the C-H stretching region (2930 - 2770 cm^-1^) from a selected point in the cell cluster at 1.7-minute time intervals for a total duration of ∼120 minutes (17 minutes for the BS and 34 minutes for LG, HG, and KCl stimulation). The raw spectra of cells and medium are reported respectively in **Supplementary Fig. 5a** and **5b**. The variations of the phospholipid absorption band at 2852 cm^-1^ during the GSIS process can be visualized in **Fig. 2e**, where the NOAS is plotted. In the LG phase (**Fig. 2e (i)**), spectra overlapped over time, suggesting that there was no significant change in MP. When HG was added to the medium (**Fig. 2e (ii)**), the NOAS of the cells at 2852 cm^-1^ increased by 24%, alluding to depolarization of the membrane and the consequent release of insulin. After the initial trigger, we observed a decrease of the NOAS of 33%, which is attributed to a return to resting state of the MP. Finally, in the last phase of the GSIS, when KCl is added (**Fig. 2e (iii)**), the NOAS increased by 60%. To summarize, **Fig. 2e (iv)** compares the mean spectra of the three different conditions (LG, HG, KCl), clearly showing how the NOAS of the phospholipid band increased upon glucose and KCl stimulation. The AUC of the NOAS at the 2852 cm^-1^ lipid absorption band as well as the relative change (% ΔNOAS/NOAS) of the spectra over time, reported respectively in **Fig. 2f** and **Supplementary Fig. 5c**, confirmed the significant change in the cells’ spectra upon HG and KCl stimulation, while the spectra of the medium was stable overtime. In detail, the AUC of the cells increased by 28% during the HG phase and by 75% during the stimulation with KCl. These observations are in agreement with the (biphasic) insulin secretion characteristics described in literature.^26^ As a control experiment, INS1 cells kept in a medium with constant amount of metabolites were spectrally monitored for the same amount of time (∼120 minutes) as stimulated INS1 cells in **Fig.2**. The raw spectra of cells and medium are reported in **Supplementary Fig. 6a, b**. As expected, the mean average spectra (**Supplementary Fig. 6c**), the line plot showing the relative change of these spectra (**Supplementary Fig. 6d**) and the line plot of their AUC (**Supplementary Fig. 6e**) did not show any changes over time. The control experiment conducted on unstimulated INS1 cells supports the hypothesis that MP changes are caused by glucose and KCl uptake that induces secretion of insulin.

In a second experiment, we confirmed our findings by monitoring the GSIS process in MIN6 cells. As with INS1 cells, spectra and images were acquired in the C-H stretching region at different stages of the insulin secretion process. **Supplementary Fig. 7a** shows the cluster of cells and medium from where spectra were acquired over time (**Supplementary Fig. 7b**). The spectra of the cells show an increase of signal after addition of glucose and KCl, as indicated also by the line plot of the AUC of the band at 2852 cm^-1^ (**Supplementary Fig. 7c**). The AUC increased by 39% and 35% during the depolarization phases corresponding to the addition of HG and KCl, respectively. The detection of depolarization during the GSIS of MIN6 was further confirmed by the analysis of the relative change of the OAC. **Supplementary Fig. 7d** illustrates representative micrographs acquired at 2852 cm^-1^ at different phases of the process, and **Supplementary Fig. 7e** summarized the change of contrast obtained from the micrographs. As for the spectral analysis, the relative change of OAC obtained from the images also exhibited voltage peaks when glucose and KCl were added. To confirm that the changes in the OA signals were caused by depolarization induced by insulin secretion, we performed control experiments on MIN6 cells, which were spectrally analyzed for 120 minutes in a medium to which no glucose or KCl had been added, similarly to what previously reported for INS1 in **Supplementary Fig. 6**. **Supplementary Fig. 8a, b** show raw spectra of MIN6 cells and medium acquired in the C-H stretching region (2930 – 2780 cm^-1^). As can be seen in **Supplementary Fig. 8c**, the NOAS obtained at the three time points corresponding to the three phases of the GSIS protocol did not change during the measurement. Indeed, the line plots of the relative change of NOAS and of the AUC of the spectra remained flat and stable throughout the measurements for both cells and medium (see **Supplementary Fig. 8d, e**). The results summarized in **Supplementary Fig. 8** are in line with the results obtained for INS1 cells (**Supplementary Fig. 6**). We further corroborated our observations by analyzing changes in OAC and NOAS in fixed MIN6 cells (**Supplementary Fig. 9**) exposed to LG, HG and KCl. Thus, we excluded that the variation of the OAS was caused by the alteration in density of the medium, which would affect the Grüneisen parameter and consequently the OAS. As shown in **Supplementary Fig. 9b,** the NOAS acquired at a selected point in a cluster of cells and the surrounding medium do not show significant changes overtime although the fixed cells were treated following the GSIS protocol (**Supplementary Fig. 9a**). The line plot of the AUC of the spectra was flat (**Supplementary Fig. 9c**), and the same lack of significant variation of OAS was also observable when analyzing the relative changes of OAC in **Supplementary Fig. 9d, e**. To further validate our results using MIRACLES, the cell media was collected at the end of each stage, and the insulin concentration was quantified using ELISA. The results are shown in **Supplementary Fig. 10**, the measured insulin levels increased from ∼2 mg/mL for LG, to ∼4 mg/mL for HG, up to ∼14 mg/mL for KCl, in agreement with the values measured by MIRACLES. The increased quantification of insulin by the ELISA confirmed the functionality of the β-cells and that the changes in OA signal observed in the stimulated experiments were caused by the cellular response (depolarization) to the stimuli.

## Discussion

Here, we introduced MIRACLES, an optoacoustic vibrational method that enables sensitive and specific detection of conformational changes within the cellular phospholipid bilayer upon MP variations. Using MIRACLES, as confirmed with standard MP assays, we demonstrated monitoring of MP in a label-free and non-destructive manner directly in living cells and applied to assess cellular functionality in β-cells upon release of insulin.

Unlike conventional MP monitoring methods, which are invasive or rely on exogenous fluorescent labels, and emerging label-free alternatives that suffer from weak signals, MIRACLES enables for the first time label-free detection of MP with high sensitivity. Indeed, while techniques such as hSRS have achieved high imaging speeds at subcellular resolution, SNR is still low (3.6) even when changes in MP raise to ∼100 mV.^15^ Additionally, hSRS was only demonstrated relying on patch-clamp stimulation to trigger changes in MP. Consequently, it remains uncertain whether hSRS can effectively capture MP changes occurring under native physiological conditions without the use of patch clamp. Moreover, MIRACLES enables monitoring of MP dynamics intrinsically, without disrupting the plasma membrane.

MIRACLES enables the mapping of spatiotemporal MP variations across cell populations, providing a more comprehensive view than single-cell sensing methods such as patch clamp, while still allowing functional assessment at the single-cell level to reveal cellular heterogeneity. Moreover, going beyond previous vibrational spectroscopy studies that have detected VSE,^14,27,28^ a critical insight of our work was clarifying that the molecular mechanism associated with MP and detectable by MIRACLES is the conformational change of phospholipids, which can be detected by a change in intensity at 2852 cm^-1^, representative band of CH_2_ symmetric stretching. Overall, our findings support the conclusion that MP changes are associated with phospholipid reorientation, as evidenced by the linear correlation between OAS and depolarization intensity. Moreover, our observations align with earlier reports from optoacoustic microscopy techniques operating in the visible range,^12^ and from polarized modulation infrared reflection spectroscopy (PM-IRRAS), which demonstrated that phospholipid headgroup and acyl-chain orientational changes are reflected in the magnitude of the applied electric field.^19^ The correlation between the amplitude changes of the electric field and the lipid orientation directly supports our hypothesis, further validated by standard MP assays, to sense the CH_2_ band associated to phospholipid as reporter of membrane-associated electrical states.

Beyond basic demonstration, we applied MIRACLES to monitor GSIS in pancreatic β-cells. In diabetes research GSIS is the gold standard to assess β-cell functionality, whose decompensation is an early biomarker of type 2 diabetes, and it is a key test to evaluate disease progression and the efficacy of therapies for restoring β-cell secretory function. GSIS quantifies the release of insulin using the Enzyme-Linked Immunosorbent Assay (ELISA), which is a highly sensitive method. However, the ELISA assay measures the average release of insulin from an unspecific cell population, and it is unable to provide insights about individual metabolic events.^7,29^ Alternatively, intracellular Ca^2+^ imaging can detect the cytosolic Ca^2+^ transients that precede insulin exocytosis,^7^ but requires exogenous labelling and can lead to phototoxic effects. Our findings link OAS signals to insulin release during depolarization, establishing MIRACLES as a robust method for assessing β-cell functionality.

Further developments of MIRACLES target the improvement of imaging speed, currently limited to one frame or one spectrum per minute. For example, MIRACLES could be incorporated into wide field modalities such as, for instance, optothermal imaging, where molecular contrast images can be acquired at larger field-of-views and higher resolution.^31–35^ Advances in imaging speed would allow extending the working applications from the static resting membrane potential to the fast transient depolarization underlying action potentials, which occur orders of magnitude faster than the current measurement rate. For example, by achieving higher frame rates, the label-free monitoring of action potential propagation across larger cell populations would become possible, enabling the study of β-cell electrical coordination without VSDs or genetic reporters. Moreover, achieving subcellular spatial resolution would further open access to organelle-level membrane potential dynamics such as mitochondrial membrane potential, providing a label-free alternative to intracellular voltage measurements.

In summary, we introduced MIRACLES as a novel label-free, non-destructive modality for mapping spatiotemporal MP dynamics in ensemble cell populations. By sensing MP induced conformational changes in the phospholipid bilayer, MIRACLES circumvents the need for invasive electrophysiological techniques, enabling the interrogation of cellular activity in native physiological states. Together, these results position MIRACLES as a versatile platform for functional phenotyping, with particular promise for β-cell assessment in drug discovery and diabetes research, and broader potential for probing other fundamental electrophysiological processes.

## Methods

### System description

MIRACLES was implemented using a tunable pulsed quantum cascade laser (QCL; MIRcat, Daylight Solutions) as the excitation source. The laser was operated across 3.41–3.61 μm (2932–2770 cm^-1^) and 5.75–11.11 μm (1739–900 cm^-1^) with a linewidth below 1 cm⁻¹, delivering 20 ns pulses at a 100 kHz repetition rate. The beam was focused into the sample using a reflective objective (36×, 0.5 NA, Newport), and the sample was mounted on a custom holder incorporating a mid-infrared–transparent ZnS or ZnSe window. OAS generated at the focal plane were detected in transmission mode with a focused ultrasound transducer (20–25 MHz central frequency; Imasonic/Sonaxis) aligned coaxially with the optical axis. Acoustic coupling was provided by the cell culture medium, ensuring efficient propagation of the OAS to the detector. The signals were amplified with a low-noise preamplifier, filtered, and digitized using a high-speed data acquisition system. The sample was raster-scanned on a motorized translation stage, enabling reconstruction of mid-infrared absorption maps. The lateral resolution of the system was experimentally determined to be approximately 5.3 μm, allowing label-free imaging of biomolecular distributions based on their vibrational absorption in the mid-infrared spectrum.

A high-precision peristaltic pump (Masterflex) was integrated with the MIRACLES setup to enable controlled exchange of cell medium during live-cell measurements. The closed-loop system contained inlet and outlet tubes connected to a modified glass reservoir, maintaining constant medium volume in the MIR dish. Flow rates were adjustable from 0.28 to 1700 mL/min depending on tubing diameter, with 10 mL/min typically used to ensure efficient circulation while minimizing shear stress. The reservoir design allowed direct addition of chemical solution to the circulating medium, enabling real-time stimulation during imaging.

### Cell culture

To evaluate the ability of MIRACLES in detecting MP changes we used non-excitable cells, such as HEK cells (HEK293, ATCC: CRL-1573), and HeLa cells (ATCC: CCL-2). HEK cells were cultured in minimum essential medium (MEM, Gibco, Fisher Scientific, USA), supplemented with 10% fetal bovine serum (FBS, Merck, DE) and 1% penicillin/streptomycin (pen/strep, Gibco, Fisher Scientific, USA). HeLa cells were cultured in Dulbecco’s modified Eagle’s medium (DMEM, Merck, DE), supplemented by 10% FBS, and 1% pen/strep. To assess changes in MP during the GSIS process, insulinoma β-cells, such as MIN6 (Merck, DE) and INS1 (INS1 832/13, Merck, DE) were used. MIN6 were grown in DMEM, supplemented with 10% FBS, 1% GlutaMAX (Gibco, Fisher Scientific, USA), 1% pen/strep, and 50 μM of 2-mercaptoethanol (Gibco, Fisher Scientific, USA). INS1 were cultured in Roswell Park memorial institute 1640 medium (RPMI 1640, Merck, DE) supplemented with 10% FBS, 1% pen/strep, 1% glutamine (Gibco, Fisher Scientific, USA), 25 mM of HEPES (Gibco, Fisher Scientific, USA), 1 mM sodium pyruvate, 50 μM of 2-mercaptoethanol (Gibco, Fisher Scientific, USA). All cells were grown in a humidified incubator at 37°C, 5% CO_2_, and were passed upon trypsin (Gibco, Fisher Scientific, USA) digestion every four days to avoid reaching confluence.

### Membrane potential stimulation

To depolarize and hyperpolarize HEK and HeLa cells (see **Fig. 1**, **Supplementary Fig. 1, 2 and 4**), cell medium containing different concentration of KCl was infused into the cells using the peristaltic pump. The initial concentration of 5.4 mM potassium chloride, which is usually present in the medium, was taken into account when varying the MP voltage values.

### Validation of cellular membrane potential

To verify that the change in OA signal caused by stimulation with different concentrations KCl caused a local change in MP, HEK cells exposed to cell medium with different concentrations of KCl were analyzed using a cellular MP assay kit (Abcam, UK). HEK cells were seeded on a poly-L-lysine coated 96-well plate; MEM medium was exchanged with a dye loading solution containing the MP sensor (fluorometric-red) and the Hank’s balanced salt solution enriched with 20 mM of Hepes buffer (HHBS). After an incubation period of 1 hour, HEK cells were exposed to different concentrations of KCl, generating a Δ potential of 0, 15, 30, 45, 60, 75, 90 mV. The red fluorescence emission of the MP dye was measured using a multi-well spectrometer at excitation and emission wavelength of 620/650 nm. To corroborate the fluorescence intensity with the OAS, HEK cells derived from the same batch were stimulated with the same concentrations of KCl and their MP was analyzed using MIRACLES (see. **Fig. 1h**).

### Glucose stimulated insulin secretion

Insulin secretion was induced following the GSIS protocol reported by Y. Zhu *et al*.^25^ INS1 and MIN6 cells were washed and placed in freshly prepared Krebs–Ringer bicarbonate HEPES (KRB) buffer containing 25.8 mM of sodium chloride (NaCl), 0.96 mM of KCl, 0.24 mM of potassium phosphate monobasic (KH_2_PO_4_), 0.24 mM of magnesium sulfate monohydrate (MgSO_4_ – H_2_O), 0.4 mM of calcium chloride (CaCl_2_), 4.8 mM of sodium bicarbonate (NaHCO_3_), and 0.1% fatty acid–free bovine serum albumin (BSA). INS1 and MIN6 cells were measured by MIRACLES for 10 minutes in the KBR buffer to determine the baseline signal (BS). Then, the cells were exposed to KBR containing 2.8 mM glucose (LG) for ∼30 minutes to establish the basal insulin secretion level. In the next step, the amount of glucose in the KBR buffer was increased to 20 mM (HG) and cells were monitored for another ∼30 minutes while secreting insulin. In the final step, the HG-KRB was substituted with KBR supplemented with 20 mM of KCl to induce the membrane depolarization.

### Image processing

Images were acquired using MIRACLES at a step size of 5 µm with a field of view of 800 x 800 μm (**Fig. 1c**), 300 x 300 μm (**Fig. 1g**), 80 x 80 μm (**Supplementary Fig. 2c**), 200 x 200 μm (**Supplementary Fig. 4a, c**), 250 × 250 µm (**Fig. 2b, Supplementary Fig. 7a, d, Supplementary Fig. 9a**), and 100 × 100 µm (**Supplementary Fig. 9d**). Segmentation of cellular and medium regions was performed using the Kittler thresholding method.^30^ Mean signal intensities were extracted separately from the cell and medium regions, and relative change in OAC (% ΔOAC/OAC) was calculated from these values. For image display purposes only, images were normalized by subtracting the minimum pixel value to enhance visual contrast. All image processing and display procedures were carried out using MATLAB (MathWorks) and ImageJ (NIH).

### Analysis of spectra

MIRACLES micrographs were acquired, and individual points were selected for spectral measurements from cells and the surrounding medium. OA spectra were recorded over the range of 2932–2780 cm^-1^ with a step size of 4 cm^-1^, yielding 39 spectral points per location. To analyze the spectra, each was normalized around 2852 cm^-1^ to generate NOAS. The NOAS intensity around the 2852 cm^-1^ lipid absorption band was extracted, and % ΔNOAS/NOAS were calculated by comparing values across time or between conditions to the initial reference value. In addition, the AUC of the NOAS was computed. All spectral analyses were performed using MATLAB (MathWorks).

## Notes

OAS: Optoacoustic Signal

NOAS: Normalized Optoacoustic Signal

OAC: Optoacoustic Contrast

### Acknowledgements

The research leading to these results has received funding from the Deutsche Forschungsgemeinschaft (DFG) - 455422993 as part of the Research Unit FOR 5298 (iMAGO, subproject TP2, GZ: NT 3/32-1; and subproject TP3, GZ: PL825/3-1) and from the Helmholtz Munich Innovation & Translation Call (OPTO-G).

## Author contributions

F.G. and J.Q. designed and performed all the imaging/spectroscopy experiments, processed the results, prepared figure panels and wrote the paper. A.A. built and characterized the MIRACLES system and performed proof-of-concept experiments. I.B. and H.L. provided support with biological knowledge about pancreatic cells. V.N. provided support on optical microscopy. M.A.P. developed the concept, designed experiments, supervised the whole study and wrote the manuscript. All authors edited the paper.

## Competing interests

M.A.P. is a founder and equity owner of sThesis GmbH. V.N. is a founder and equity owner of Maurus OY, sThesis GmbH, Spear UG, Biosense Innovations P.C. and I3 Inc. V.N. and M.A.P. are inventors on a provisional patent application related to mid-IR optoacoustic microscopy (US-2020355604-A1, US Patent, 2018). The other authors declare no competing interests.

We thank S. Lee for assisting with editing of the paper.

We also thank Dr. Ciro Salinno for providing initial support in handling pancreatic beta cells.

---

**Supplementary Fig. 1.**
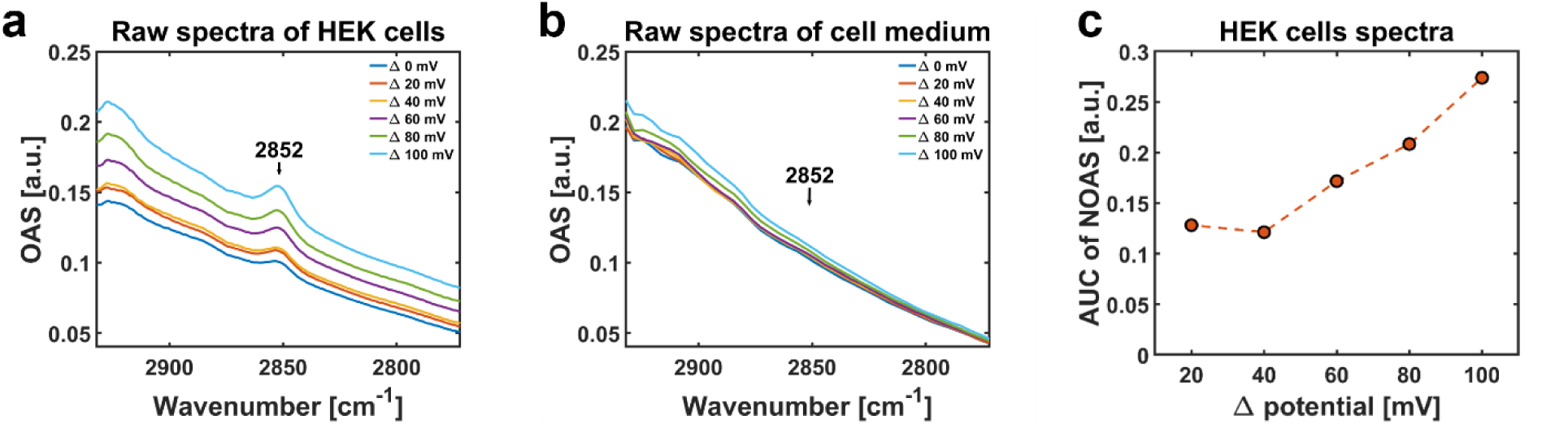
Spectra of HEK cells during the depolarization process. **a)** Raw spectra of HEK cells and **b)** cell medium acquired during the depolarization process shown in Fig. 1c**-g**. HEK cells were stimulated with increasing concentration of potassium chloride (KCl) and their absorption band at 2852 cm^-1^ was increasing progressively. Contrary, no significant changes were observed in the spectra of the cell medium acquired overtime. **c)** Area under curve (AUC) of the spectra in supplementary Fig. 1 a, b and Fig. 1d. OAS: optoacoustic signal. NOAS: normalized optoacousticsignal.

**Supplementary Fig. 2.**
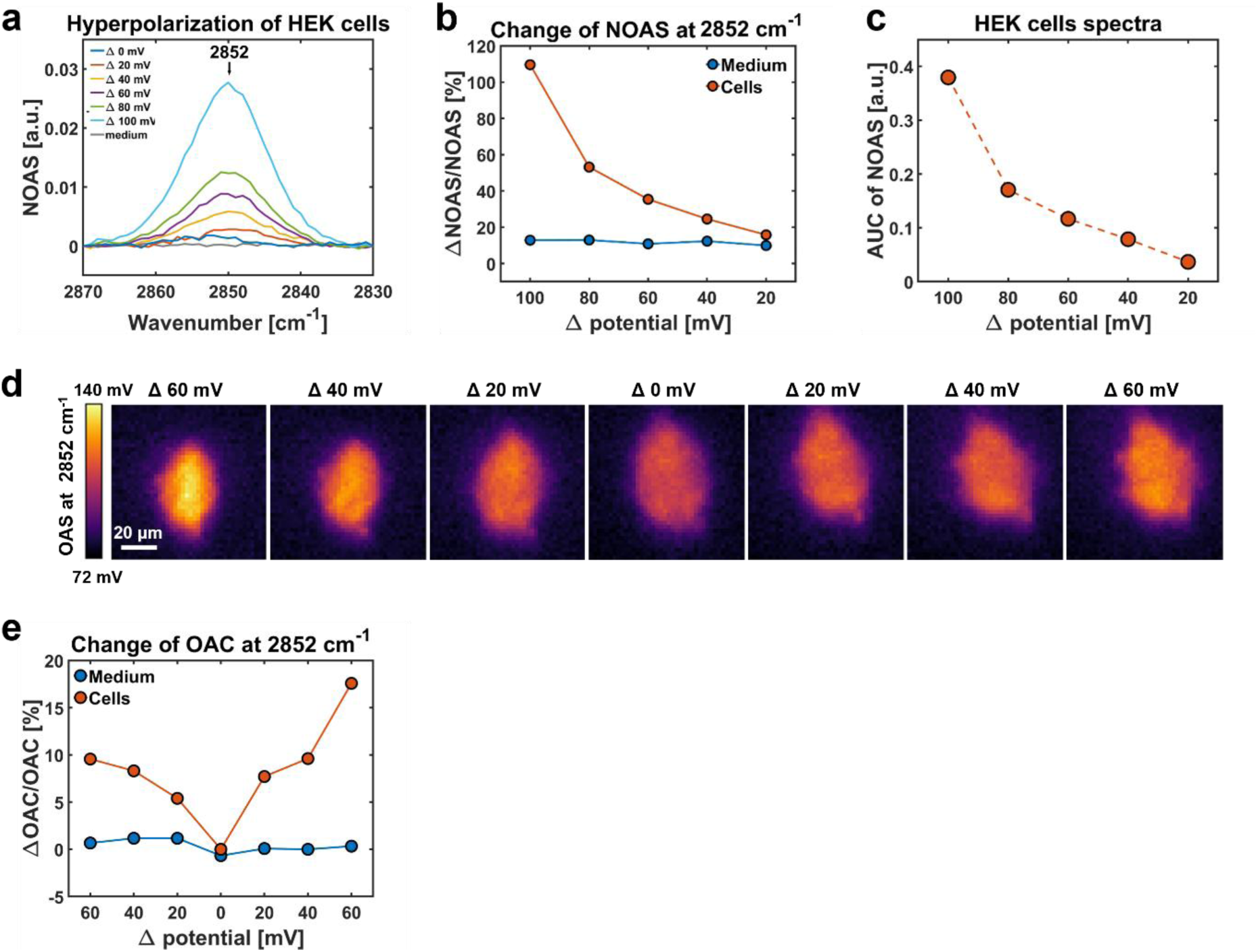
Hyperpolarization of human embryonic kidney (HEK) cells. HEK cells were hyperpolarized through stimulation with progressively low concentrations of potassium chloride (KCl). **a)** Normalized mean spectra of HEK cells and cell medium. The absorption peak at 2852 cm^-1^ decreased while the concentration of KCl inside the medium was diluted. **b)** Line plots showing the relative change of the normalized optoacoustic signal (ΔNOAS/NOAS) at 2852 cm^-1^ for HEK cells and cell medium during the hyperpolarization process. **c)** Area under curve (AUC) of the normalized spectra in (a). **d)** Micrographs acquired at 2852 cm^-1^ showing the changes of optoacoustic contrast (OAC) in HEK cells first subjected to hyperpolarization and then to depolarization of the membrane. **e)** Line plots showing the relative change of the optoacoustic contrast (ΔOAC/OAC) obtained for HEK cells and cell medium during the hyperpolarization and depolarization of the cell membrane.

**Supplementary Fig. 3.**
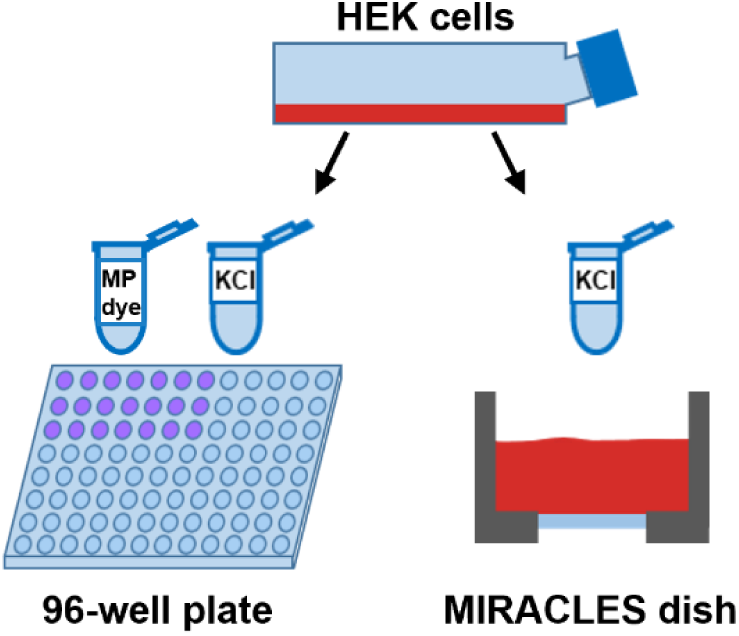
Assessment of membrane potential (MP) changes via MP assay kit. Changes of MP were evaluated using mid-infrared assessment of conformation in lipids by ensemble sensing (MIRACLES) and a membrane potential assay kit on HEK cells derived from the same passage. The fluorescent MP dye and potassium chloride (KCl) were added to HEK cells, and the MP was assessed by measuring the fluorescence emission at 650 nm using a microplate reader. MP changes were assessed by MIRACLES stimulating HEK cells with the same concentrations of KCl that were used to evaluate MP via MP assay kit.

**Supplementary Fig. 4.**
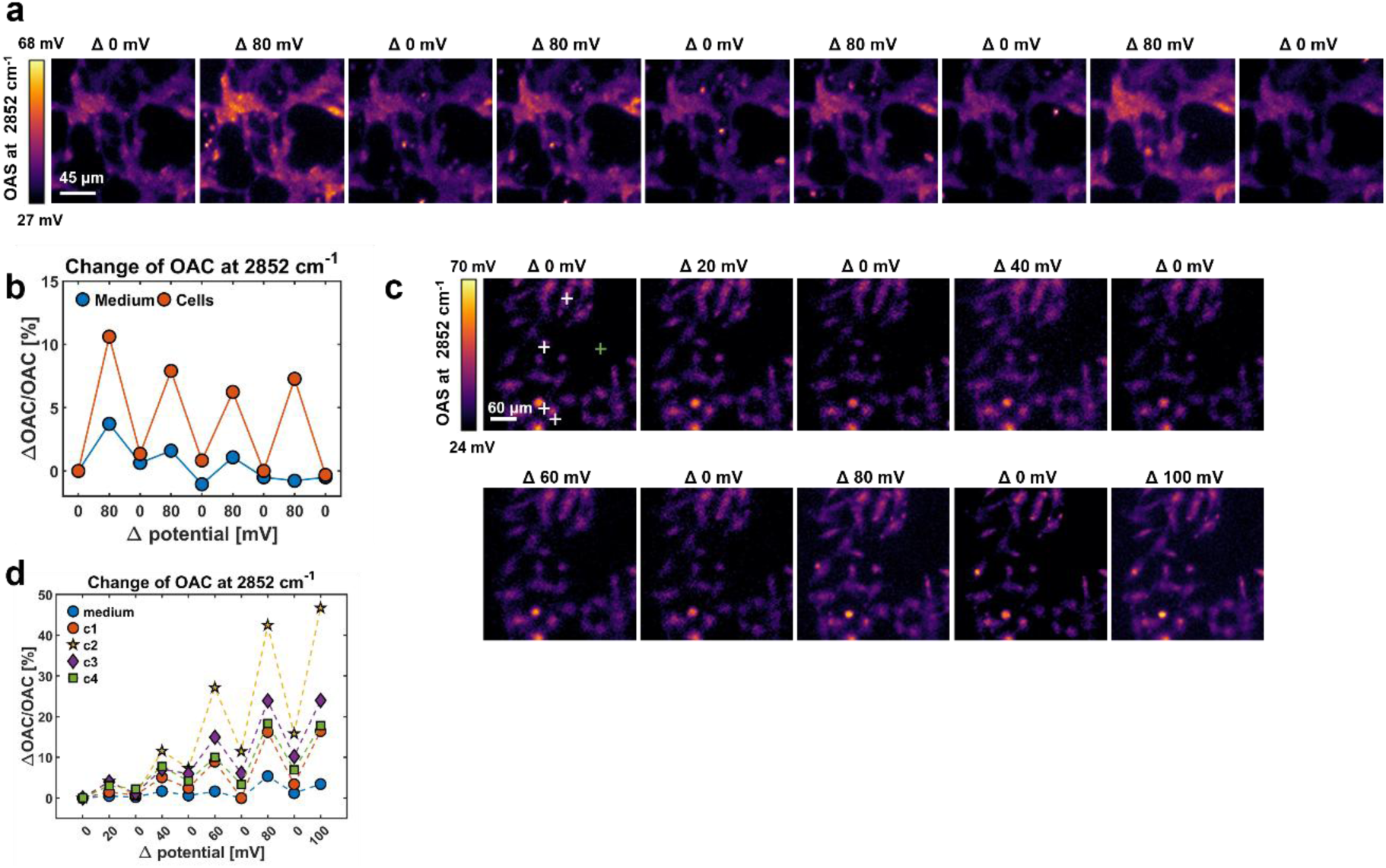
Modulation of membrane potential (MP). **a)** Micrographs of HEK cells acquired at 2852 cm^-1^ during a repetitive change of potassium chloride (KCl) concentration in the cell medium. **b)** Line plots showing the relative change of the optoacoustic contrast (ΔOAC/OAC) at 2852 cm^-1^ for HEK cells and cell medium in (a). **c)** Micrographs of HeLa cells acquired at 2852 cm^-1^ during modulation of MP alternating between a depolarization phase (increasing the concentration of KCl) to a resting potential state (Δ 0 mV). **d)** Line plots showing the relative change of OAC in four cells (white crosses) and in the cell medium (green cross). The OAC increased at increasing potential (ΔV) and decreased to lower values when the cells were in a resting potential state (Δ 0 mV). OA: optoacoustic signal.

**Supplementary Fig. 5:**
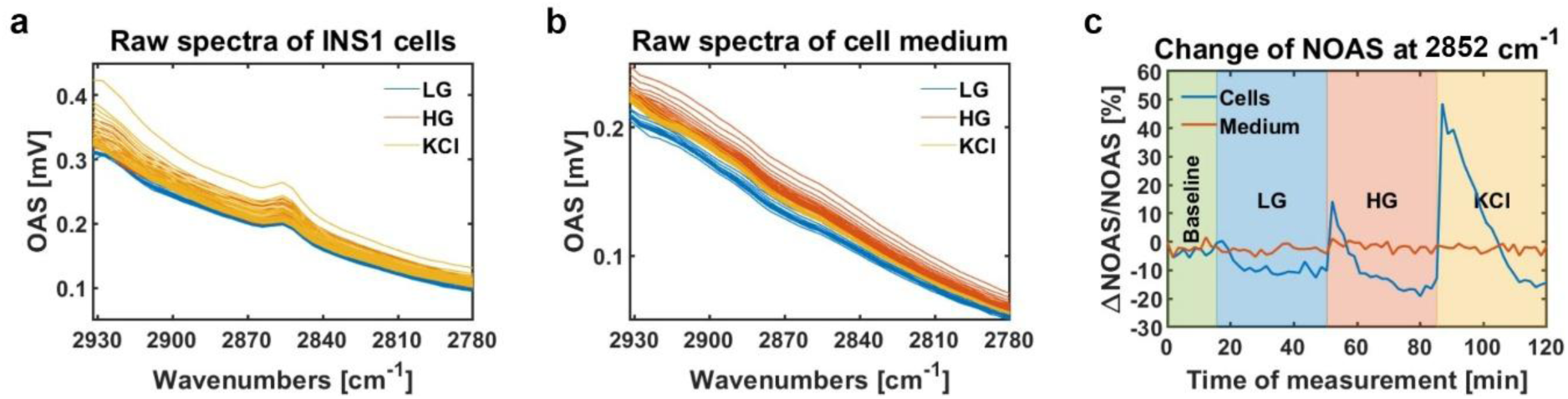
Spectra of rat insulinoma cells (INS1) during the glucose stimulated insulin secretion (GSIS) process. **a)** Raw spectra of INS1 cells and **(b)** cell medium acquired during the GSIS process. INS1 cells were exposed to concentrations of low glucose (LG), high glucose (HG) and potassium chloride (KCl). **c)** Line plots showing the relative change of the normalized optoacoustic signal (ΔNOAS/NOAS) at 2852 cm^-1^ for INS1 cells and medium in (a) and (b). OA: optoacoustic.

**Supplementary Fig. 6:**
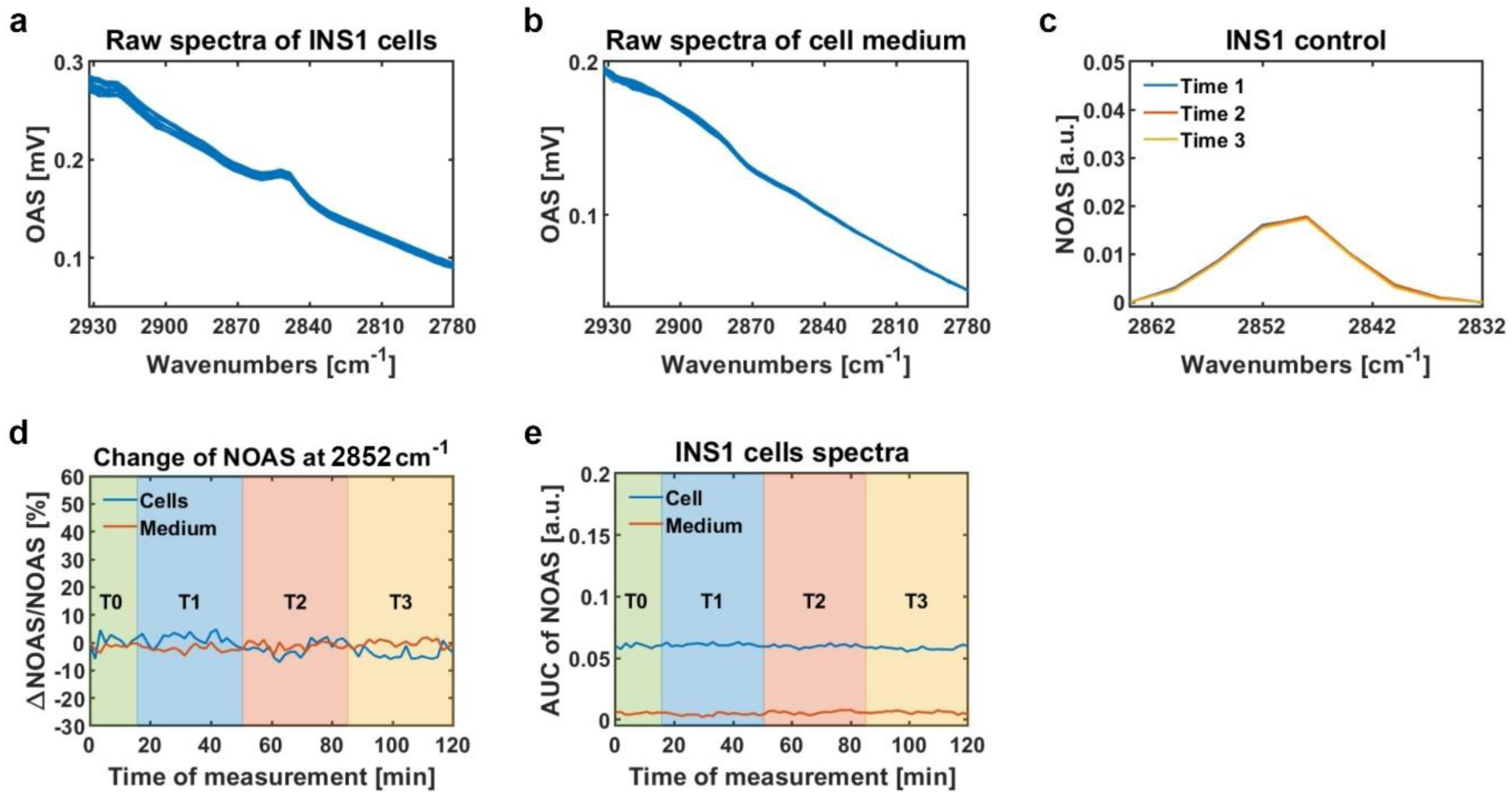
Spectra of unstimulated rat insulinoma (INS1) cells (control). **a)** Raw optoacoustic (OA) spectra acquired from single points of INS1 cells and **b)** cell medium unstimulated (control). **c)** Normalized optoacoustic spectra (NOAS) at different time points corresponding to the time points measured in the stimulated INS1 (Fig. 2c). **d)** Line plots showing the relative change of the NOAS (ΔNOAS/NOAS) at 2852 cm^-1^ obtained from the spectra of INS1 cells and medium in (a) and (b). **e)** Area under the curve (AUC) of the NOAS acquired over the time of the measurement.

**Supplementary Fig. 7:**
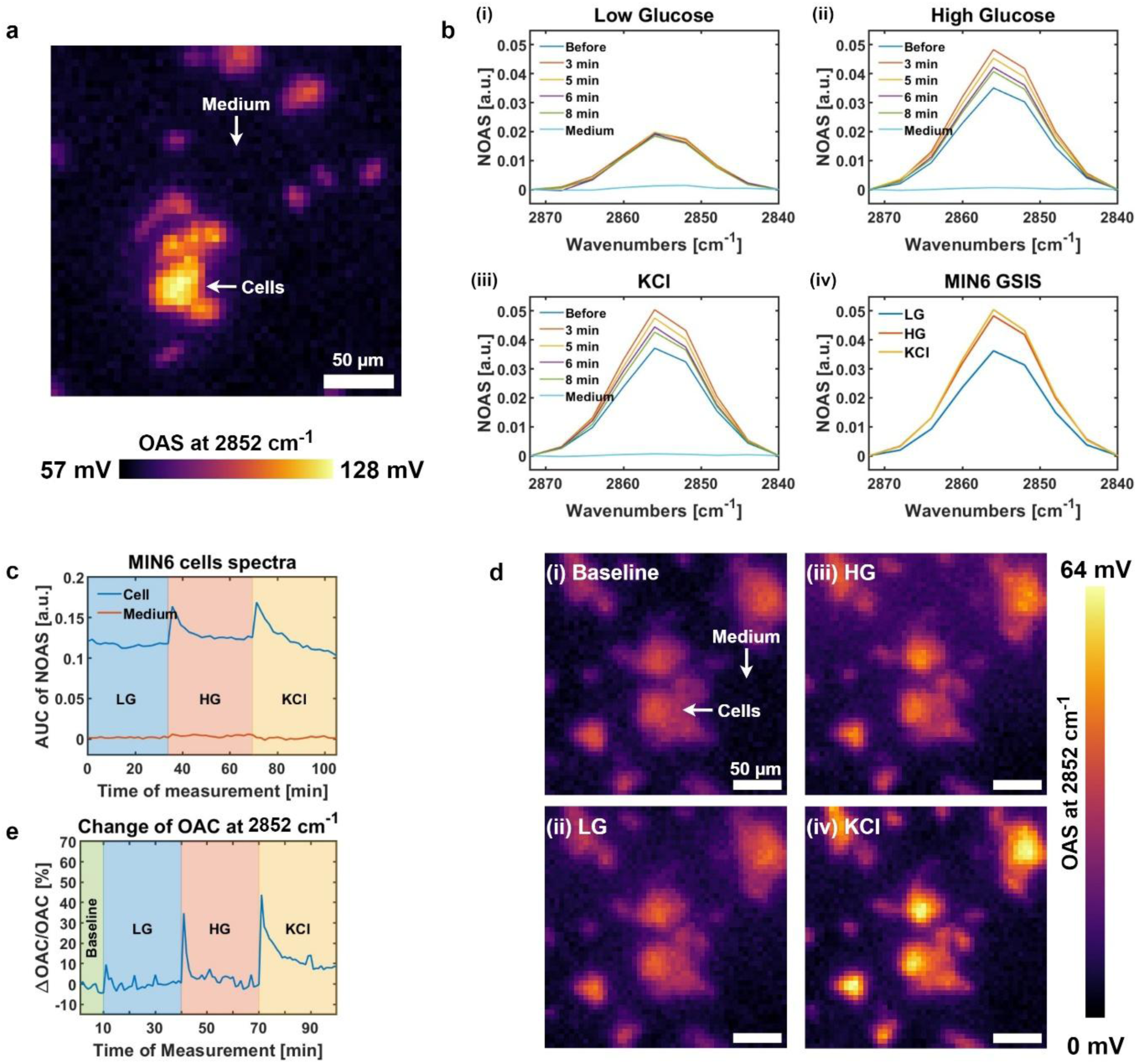
Monitoring glucose stimulated insulin secretion (GSIS) in mouse insulinoma (MIN6) cells. **a)** Micrograph of MIN6 cells acquired at 2852 cm^-1^. **b)** Normalized optoacoustic spectra (NOAS) acquired from single points in MIN6 cell and cell medium (arrows in **a)**) during the different phases of the GSIS process; (i) low glucose (LG) concentration, (ii) high glucose (HG) concentration and (iii) potassium chloride (KCl) concentration, (iv) NOAS summarizing the different phases of the GSIS process. **c)** Area under the curve (AUC) of the NOAS in (b). The increase of the AUC value after the HG and KCl phases suggests membrane depolarization. **d)** Representative micrographs of MIN6 cells acquired at 2852 cm^-1^ while changing glucose and KCl concentrations in the cell medium according to the GSIS protocol. **e)** Line plot showing the relative change of the optoacoustic contrast (ΔOAC/OAC) of cells and cell medium calculated from the micrographs in **d)**. OA: optoacoustic.

**Supplementary Fig. 8:**
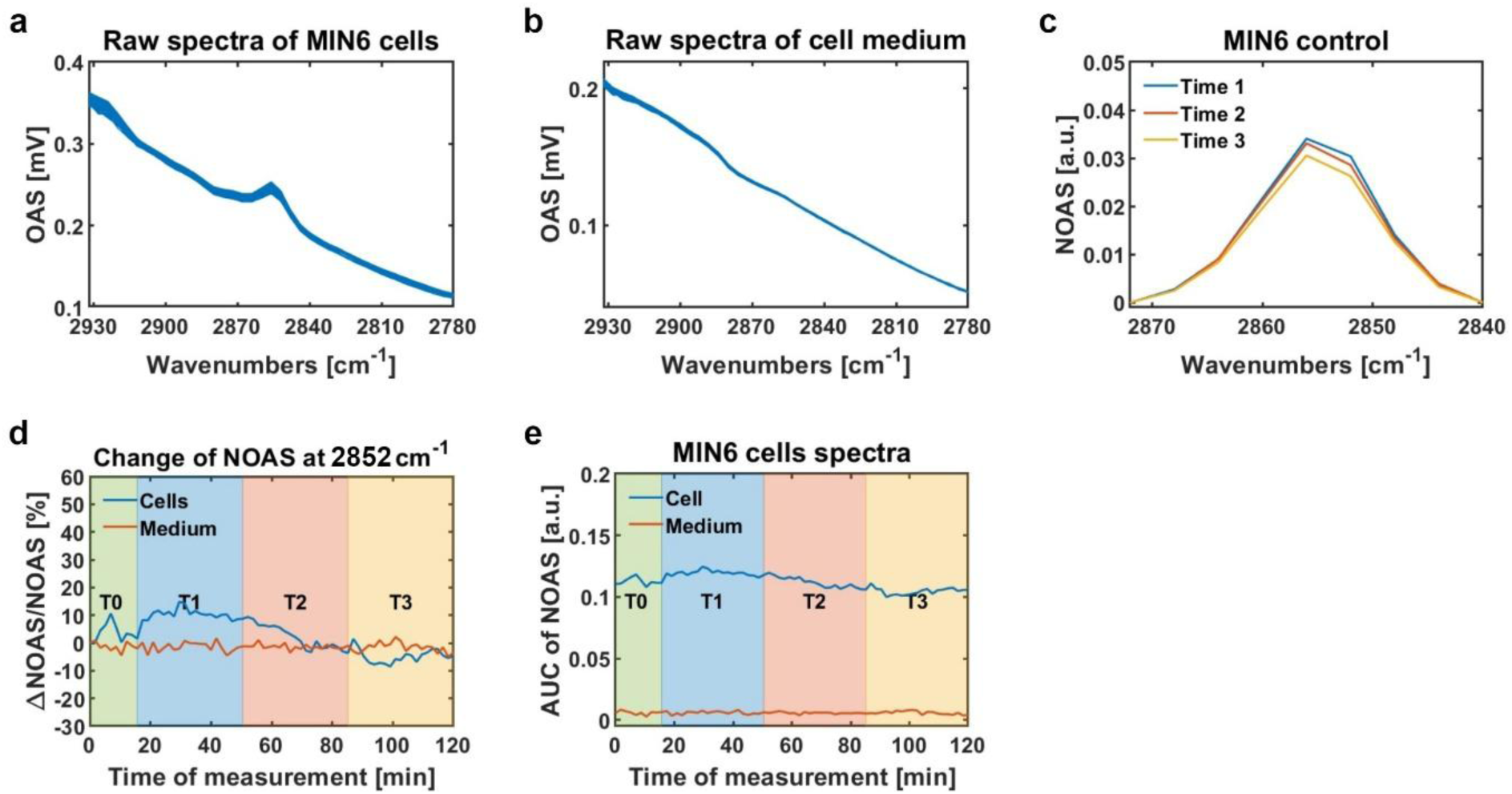
Spectra of unstimulated mouse insulinoma (MIN6) cells (control). **a)** Raw optoacoustic (OA) spectra of MIN6 cells and **b)** cell medium unstimulated (control). **c)** Normalized optoacoustic spectra (NOAS) at different time points corresponding to the time points measured in the stimulated MIN6. **d)** Line plots showing the relative change of the NOAS (ΔNOAS/NOAS) at 2852 cm^-1^ obtained from the OA spectra of MIN6 cells and medium in (a) and (b). **e)** Area under the curve (AUC) of the NOAS acquired over the time of the measurement.

**Supplementary Fig. 9:**
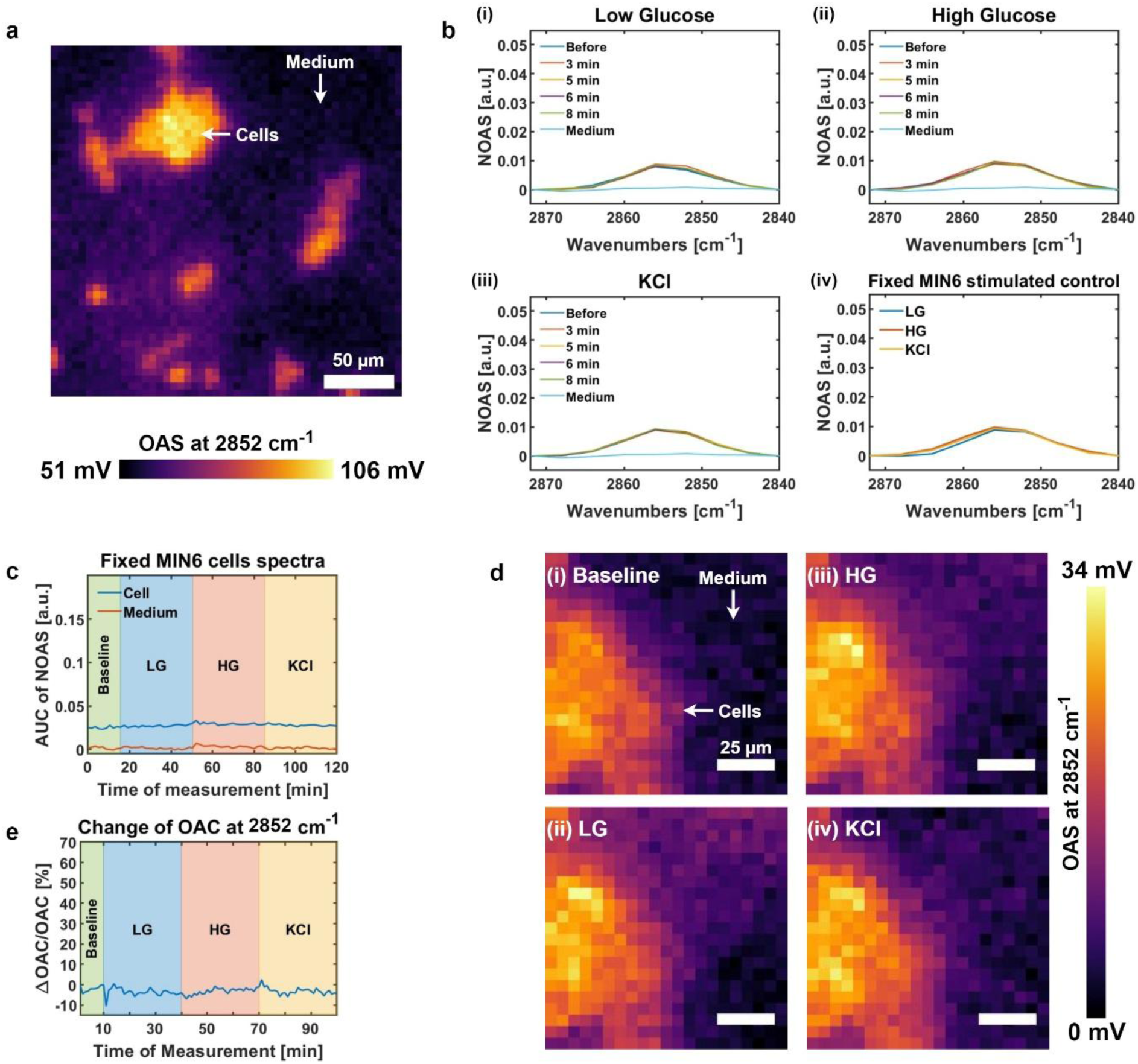
Monitoring glucose stimulated insulin secretion (GSIS) in fixed mouse insulinoma (MIN6) cells. **a)** Micrograph of fixed MIN6 cells acquired at 2852 cm^-1^. **b)** Normalized optoacoustic spectra (NOAS) acquired from single points in fixed MIN6 cell and cell medium (arrows in **a)**) during the different phases of the GSIS process; (i) low glucose (LG) concentration, (ii) high glucose (HG) concentration and (iii) potassium chloride (KCl) concentration, (iv) NOAS summarizing the different phases of the GSIS process. **c)** Area under the curve (AUC) of the spectra in (b). **d)** Representative micrographs of MIN6 cells acquired at 2852 cm^-1^ while changing glucose and KCl concentrations in the cell medium according to the GSIS protocol. **e)** Line plot showing the difference between the optoacoustic (OA) signal of cells and cell medium calculated from the micrographs in **d)**.

**Supplementary Fig. 10:**
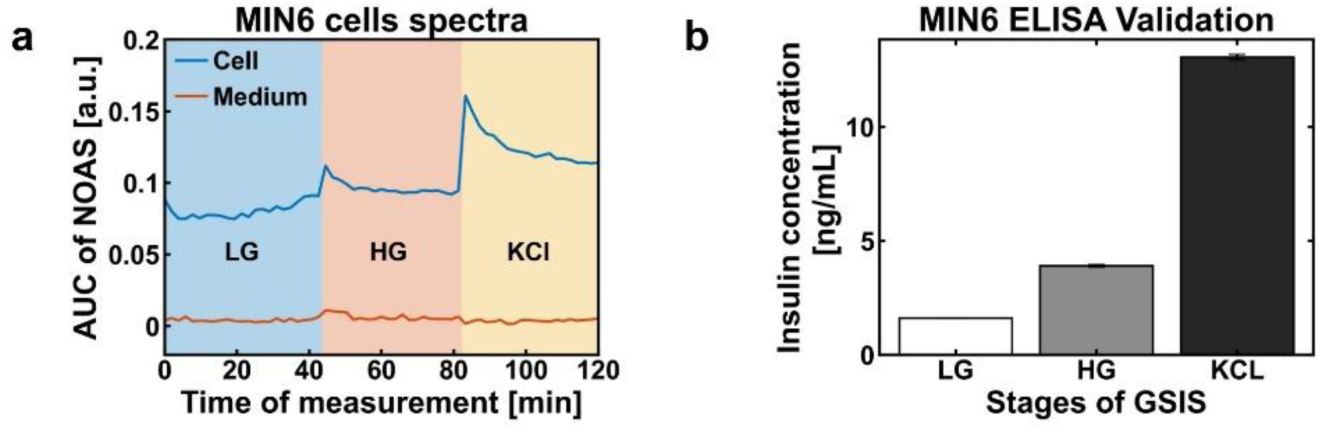
Validating glucose stimulated insulin secretion (GSIS) monitoring using Enzyme-Linked Immunosorbent Assay (ELISA). **a)** Area under the curve (AUC) of the spectra acquired with mid-infrared assessment of conformation in lipids by ensemble sensing (MIRACLES) during the glucose-stimulated insulin secretion (GSIS) process. **b)** Insulin quantification using ELISA, corresponding to each of the stages of the GSIS process.

